# Analyzing bivariate cross-trait genetic architecture in GWAS summary statistics with the BIGA cloud computing platform

**DOI:** 10.1101/2023.04.28.538585

**Authors:** Yujue Li, Fei Xue, Bingxuan Li, Yilin Yang, Zirui Fan, Juan Shu, Xiaochen Yang, Xiyao Wang, Jinjie Lin, Carlos Copana, Bingxin Zhao

## Abstract

As large-scale biobanks provide increasing access to deep phenotyping and genomic data, genome-wide association studies (GWAS) are rapidly uncovering the genetic architecture behind various complex traits and diseases. GWAS publications typically make their summary-level data (GWAS summary statistics) publicly available, enabling further exploration of genetic overlaps between phenotypes gathered from different studies and cohorts. However, systematically analyzing high-dimensional GWAS summary statistics for thousands of phenotypes can be both logistically challenging and computationally demanding. In this paper, we introduce BIGA (https://bigagwas.org/), a website that aims to offer unified data analysis pipelines and processed data resources for cross-trait genetic architecture analyses using GWAS summary statistics. We have developed a framework to implement statistical genetics tools on a cloud computing platform, combined with extensive curated GWAS data resources. Through BIGA, users can upload data, submit jobs, and share results, providing the research community with a convenient tool for consolidating GWAS data and generating new insights.

---

The rapid development of biobank-scale biomedical databases, encompassing phenotyping and genomic data, has occurred globally^1^. Numerous genome-wide association studies (GWAS) have been conducted to determine the genetic architecture underlying a wide range of complex traits and clinical outcomes, with the aim of improving disease prevention and treatment^2^. Publicly available GWAS summary-level data (or GWAS summary statistics) encompass thousands of phenotypes^3-8^. These summary statistics, derived from large-scale studies, provide valuable opportunities for in-depth investigations into genetic overlaps and shared architectures between phenotypes across studies and cohorts. Various statistical genetic tools have been developed to analyze GWAS summary statistics and examine the shared genetic components between pairs of phenotypes, such as LDSC^9^, LAVA^10^, SumHer^11^, and Popcorn^12^. These methods offer insights into genetic links from various perspectives and have been widely applied to clinical biomarkers and outcomes^13,14^.

However, implementing and batch-running these tools often requires robust computing and data infrastructure, which may not always be available to all researchers. Consequently, systematic bivariate cross-trait analyses using massive GWAS summary statistics for thousands of phenotypes can be logistically and computationally challenging. As more complex and deep phenotyping data are obtained from biobanks^15^, addressing these limitations becomes increasingly urgent. For example, the UK Biobank (UKB) imaging study^16^ collected multimodal brain imaging data, generating over 5,000 imaging-derived phenotypes using different imaging modalities and processing pipelines^17-21^. Researchers interested in a specific disease and its genetic connections with imaging biomarkers have traditionally downloaded all the GWAS summary statistics for over 5,000 imaging biomarkers from the Oxford BIG40 Project (http://big.stats.ox.ac.uk) and the BIG-KP project (https://bigkp.org/), and run their statistical tools in local clusters, which can be inefficient. Such challenges are also present in centralized GWAS databases, such as GWAS Catalog^3^ and IEU OpenGWAS^7^, where users are expected to download and manage large datasets locally to conduct most analyses. Several online research platforms based on cloud computing have been developed, most of which focus on one database (such as the UKB study, https://ukbiobank.dnanexus.com/), univariate trait GWAS analysis (such as FUMA^22^), or single data analysis method/function (such as LD Hub^23^ and Locus Compare^24^). Developing an integrated platform for cross-trait analyses of GWAS summary data resources will make existing large-scale GWAS summary data more accessible to researchers.

To address these limitations, we developed BIGA (https://bigagwas.org/), an online cloud-based platform that offers unified data harmonization and analysis pipelines and processed data resources for cross-trait analyses using GWAS summary statistics. BIGA aims to provide various tools for quantifying cross-trait genetic architectures, such as genome-wide genetic correlation methods (e.g., LDSC^9^, Popcorn^12^, and SumHer^11^) and local genetic correlation analysis (e.g., LAVA^10^). We have also aggregated and harmonized GWAS summary statistics from various resources, including the GWAS Catalog^3^, UKB study^15^, Psychiatric Genomics Consortium^25^, FinnGen^6^, Biobank Japan^8^, CHIMGEN^26^, UKB-PPP^27^, BIG-KP^18,19,21^, and Oxford BIG40^17,20^. These curated datasets, currently including over 15,000 traits, have been integrated with multiple methods, facilitating easy online analysis for users. With our established infrastructure in place, we are committed to the continuous development and growth of BIGA, aiming to broaden its capabilities by consistently including new tools and data resources.

**Figure 1.** provides an overview of the BIGA architecture. We offer users several options for inputting GWAS summary statistics data with user-friendly features, including uploading their own data, querying data from public databases (such as the IEU OpenGWAS^7^, GWAS Catalog^3^, and Neale Lab (http://www.nealelab.is/uk-biobank), and reusing data from recent previous jobs (Supplementary Text). Users can specify the tools and job types they are interested in and submit their requests. After submission, the job request will be passed to the back-end and executed on our cloud computing platform using the specified tools and datasets. Briefly, we have developed a thorough pipeline for harmonizing user-input data, similar to procedures used in the GWAS Catalog (https://github.com/EBISPOT/gwas-sumstats-harmoniser). After harmonization, datasets will have a standard format with column names outlined in **Table S1**. Considering the specific data format needed by the user-requested analysis, we will accordingly adapt the data to fulfill these requirements and execute the analysis (**Fig. S1**) Once completed, users will receive email notification and the results will be presented to the users through the front-end interface. A quick-start tutorial and comprehensive documentation are available on our website for users.

**Fig. 1.**
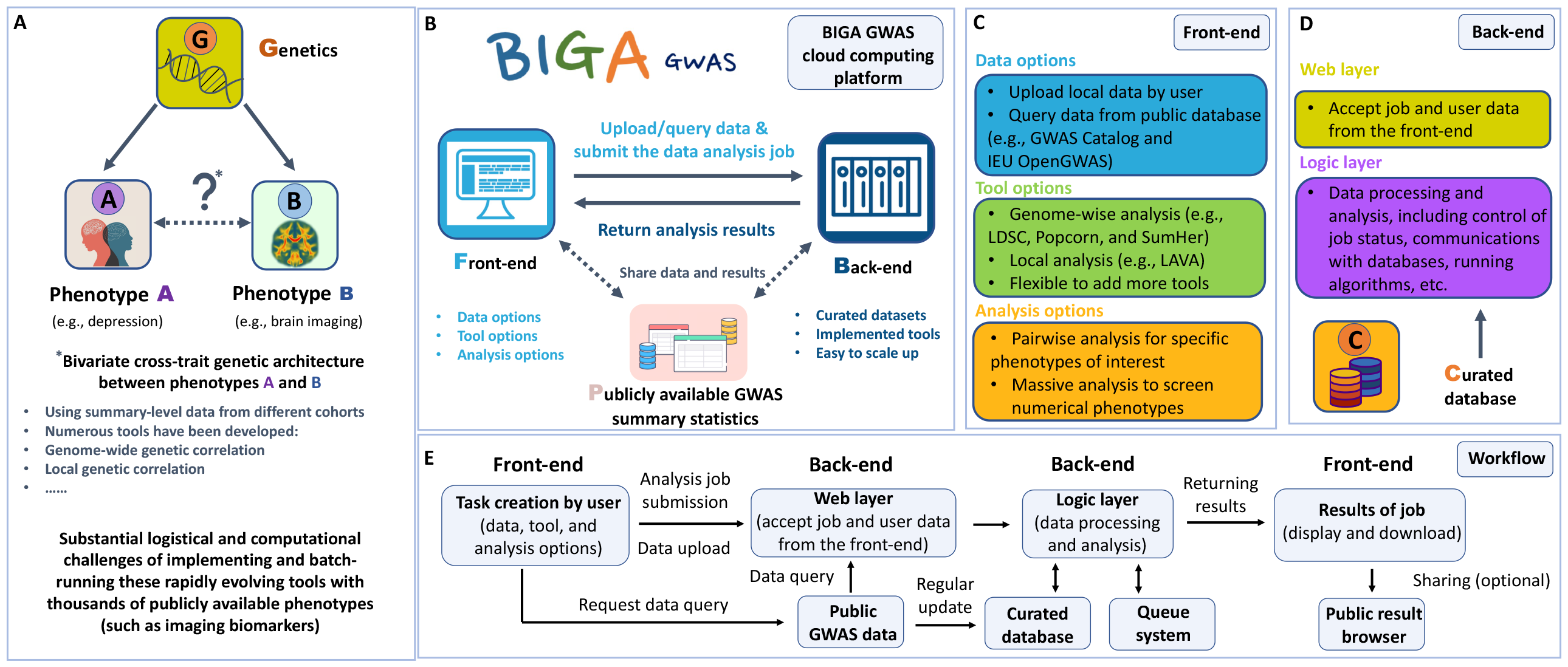
Overview of BIGA GWAS cloud computing platform. **(A)**The motivation of this project is to address the substantial logistical and computational challenges associated with implementing and batch-running the constantly evolving tools for cross-trait genetic architecture analysis. Our aim is to offer a cloud computing-based solution that can effectively overcome these challenges. **(B)** Overview of the BIGA GWAS platform. Users can easily upload or query GWAS summary statistics and submit data analysis jobs through the front-end interface. These jobs are then processed on the back-end, and the results are subsequently returned to the users. **(C)** The front-end interface of the BIGA GWAS platform offers users a comprehensive set of options to manage their data resources, choose the appropriate tools, and select the desired mode of data analysis. **(D)** Details of the back-end of the BIGA GWAS platform. **(E)** Overview of the analysis workflow.

BIGA uses a powerful and efficient computational framework for automated analysis. Every step, from the initial data input to the final results output, is organized by a standardized pipeline, offering the flexibility to incorporate new methods. For example, BIGA operates on the Django 3.2 web framework (https://www.djangoproject.com/) to accommodate various tasks and tools, and we use Redis (https://redis.io/) and Celery (https://docs.celeryq.dev) for task management and queuing system. BIGA’s computational infrastructure is efficient, currently supporting 20 concurrent user jobs running with just 128GB of RAM and 16 Intel vCPUs. Notably, cloud computing services provide a flexible management system for CPU and RAM, enabling us to easily modify our resource allocation for scaling up or down as needed. Even with only 16GB of RAM, BIGA can execute 3 jobs concurrently using our efficient configuration. We have conducted large-scale tests to validate the stability and computational efficiency of BIGA (**Figs. S2-3** and Supplementary Text).

To showcase the extensive genetic analyses that BIGA can conduct, we present a blood pressure data analysis example, aiming to explore its genetic correlation with over 15,000 complex traits and diseases curated on BIGA. We initiated the analysis by searching for blood pressure data on the IEU OpenGWAS database and used the BIGA query function to directly query systolic blood pressure^28^ summary statistics. BIGA performed harmonization and then used the harmonized data to run LDSC massive analysis, spanning over all groups of traits from European population on BIGA. As expected, at a false discovery rate 5% level, systolic blood pressure was widely associated with complex traits and diseases, such as hypertension, atrial fibrillation, stroke, brain and body imaging traits, as well as plasma proteomics (*P* range = (5.44×10^−244^, 4.00×10^−2^), **Fig. S4**). We further examined the diastolic blood pressure^28^ and found similar association patterns to systolic blood pressure (**Fig. S5**). We applied SumHer to repeat the analysis (**Fig. S6**) and observed that the results from LDSC and SumHer were generally consistent (**Fig. S7**, Pearson’s correlation = 0.9273). In addition, we performed local genetic correlation analysis using LAVA and cross-population genetic correlation using Popcorn. More details can be found in the Supplementary Text (**Tables S2-S8**). This data analysis example demonstrates that BIGA facilitates efficient analysis of extensive GWAS summary statistics with different methods.

In summary, our platform enables researchers to easily perform multiple cross-trait analyses without needing access to a local research computing cluster, implementing methods locally, or downloading large datasets. BIGA will help reduce the imbalance in the research community caused by unequal computing resources and attract a wider user base to these developed methods. The source code to build the BIGA platform will be made publicly available on GitHub. The BIGA website welcomes user feedback and requests, which aids in improving the project and implementing new tools and functions to better meet the needs of the research community.

## ADDITIONAL INFORMATION

*One supplementary pdf file and one supplementary table zip file are available*.

## ACKNOWLEDGEMENTS

We thank for the helpful conversions with Doug Speed regarding SumHer. Research reported in this publication was supported by the National Institute On Aging of the National Institutes of Health under Award Number RF1AG082938. The content is solely the responsibility of the authors and does not necessarily represent the official views of the National Institutes of Health. The study has also been partially supported by funding from the Wharton Dean’s Research Fund, Analytics at Wharton, NSF Grant DMS 2210860, and Purdue Statistics Department. We would like to thank the research computing groups at Purdue University, the Wharton School of the University of Pennsylvania, and the University of North Carolina at Chapel Hill for providing computational resources and support that have contributed to these research results. We would like to thank all the developers of the tools and methods implemented in our project. We gratefully acknowledge all the studies and databases that made GWAS summary data available and thank the individuals who represented these studies for their participation and the research teams for their work in collecting, processing, and disseminating these datasets for analysis.

## AUTHOR CONTRIBUTIONS

Y.L. and B.Z. designed the study, developed the BIGA website, and wrote the manuscript with feedback from all authors. B.L. helped with the implementation of statistical genetic methods and website functions. Y.Y., Z.F., J.S., X.Y., X.W., B.L., and C.C. processed the GWAS summary statistics, developed the curated datasets, and contributed to the development of the website. F.X. and J.L. provided feedback on the study design and website.

## CORRESPONDENCE AND REQUESTS FOR MATERIALS

should be addressed to B.Z.

## Code availability

All source code to develop the BIGA platform will be made publicly available on the BIGA GitHub repository. The statistical tools and methods implemented in the BIGA platform are also open source, and their source code has already been made available to the public by their authors. A summary of our implemented tools and data resources can be found at https://bigagwas.org/documentation.

## Data availability

GWAS summary statistics used in the BIGA platform are publicly available and can be found in several public databases, such as the

Neale Lab UK Biobank Results (http://www.nealelab.is/uk-biobank),

Psychiatric Genomics Consortium (https://pgc.unc.edu/),

IEU OpenGWAS (https://gwas.mrcieu.ac.uk/),

FinnGen (https://www.finngen.fi/en/access_results),

Biobank Japan (https://pheweb.jp/),

BIG-KP (https://bigkp.org/),

Oxford BIG40 (https://open.win.ox.ac.uk/ukbiobank/big40/),

UKB-PPP (https://metabolomips.org/ukbbpgwas/),

GWAS Catalog (https://www.ebi.ac.uk/gwas/downloads/summary-statistics), and

CHIMGEN (http://chimgen.tmu.edu.cn/en/index.php?c=article&id=2036).

