## Supplementary material for "Analyzing bivariate cross-trait genetic architecture in GWAS summary statistics with the BIGA cloud computing platform": supp_info

**This PDF file includes:**

Supplementary Text

Supplementary Figures Figs. S1-S7

Legends for Table S1 to S8

**Other Supplementary materials for this manuscript include the following:**

Table S1 to S8 (.xlsx) (available in a zip file)

### Supplementary Text

#### Data processing

BIGA has developed a unified data management system for massive analysis and pairwise analysis. We offer three options for inputting GWAS summary statistics: “Upload data”, “Query data”, and “Use previous data”. For the “Upload data” option, users will upload summary statistics that meet our column requirements (**Table S1**), specify the human genome version, and provide the ancestry of the summary statistics. The “Query data” function allows users to directly query summary statistics from external data sources, including the IEU OpenGWAS, Neale Lab, and GWAS Catalog, without the need to download and process the data themselves. The “Use previous data” function enables users to use data from recent previous jobs without repeatedly uploading or querying GWAS summary statistics.

Once the data is uploaded or queried, BIGA will conduct data harmonization for non-harmonized data. This harmonization process follows the steps used in the GWAS Catalog (https://www.ebi.ac.uk/gwas/docs/methods/summary-statistics), including four major parts: 1) mapping variant IDs to locations, either by rsID or base pair location; 2) inferring the orientation of palindromic variants; 3) variant harmonization with Ensembl VCF reference; and 4) filtering invalid records. During harmonization, we will convert each input summary statistics dataset into a standardized format within BIGA, and then format the data according to the specific requirements of the chosen method. The curated datasets available in BIGA have been harmonized following these steps except for a few datasets in Psychiatric Genomics Consortium (PGC) due to missing specific columns. Note that since data that can be queried from GWAS Catalog are already harmonized, we will skip data harmonization steps and directly transfer GWAS Catalog data into the standard format in BIGA. Through the standardized format, users can easily reuse data across different analysis methods. After the job is complete, users can download data processing log file and the harmonized data, which include a column names “hm_code”, recording the harmonization operations performed by BIGA on each variant. Our input data processing procedure is summarized in **Figure S1**. More details are available in our online tutorials at https://bigagwas.org/tutorials.

**Computational performance and stability of BIGA**

We have tried to improve the computational performance of our platform by reducing the memory usage for each job, especially to reduce the memory usage of the data harmonization part. Specifically, we process summary statistics using chunking and multiprocessing. This segmented approach allows us to process summary statistics in small memory usage, which substantially reduce CPU usage for a single operation. Moreover, by processing summary statistics for chromosomes in parallel, we accelerate the data harmonization time. For example, during data harmonization, we currently set chunk size to 200,000 to process 200,000 rows at a time, making the maximum CPU usage for a single operation is around 1GB per CPU. At the same time, we allocate 4 CPUs to process 4 chromosomes in parallel, lowering the overall memory usage for each job to around 5GB. As a comparison, the minimum memory usage for the original GWAS Catalog harmonization software (https://github.com/EBISPOT/gwas-sumstats-harmoniser) is around 28GB since they process the entire data at once.

To evaluate BIGA's computational performance, we conducted comparative tests with two major university-wide computing clusters accessible to us: the Purdue Bell and UNC Longleaf. We selected a set of traits comprising 101 regional brain volumes to perform a massive LDSC analysis, using two datasets: the "PGC (and brain disorders)" dataset with 24 traits and the "Brain and Organ Imaging Traits" dataset with 360 regional amplitude traits. To ensure a fair comparison, we standardized the CPU memory across all platforms to 16GB. The results showed that the average processing times were comparable: for the PGC dataset, BIGA took 175.49 seconds per job, Purdue Bell 171.31 seconds, and UNC Longleaf 175.12 seconds; for the amplitude traits, BIGA needed 2462.15 seconds, Purdue Bell 2383.55 seconds, and UNC Longleaf 2298.43 seconds. As illustrated in **Figure S2**, although BIGA's average execution time is slightly longer than that of the other platforms, it demonstrates greater stability during execution compared to the other platforms.

To further evaluate BIGA's stability under heavy load, we performed a stress test with 50 simultaneous submissions, including 33 pairwise analysis and 17 massive analysis jobs. These jobs encompass various user-input options, including querying data, uploading data, using previous data. This test was conducted during BIGA's initial setup, we configured the cloud server with 16GB of RAM and 8 Intel vCPUs. With this setup, our configuration supported the parallel execution of 2 LDSC or SumHer jobs alongside 1 LAVA or Popcorn job. BIGA managed to start with 3 active jobs while queuing 47 others, completing all tasks successfully without any CPU or RAM deficits (**Fig. S3**).

**Blood pressure data analysis with BIGA**

We performed data analysis of blood pressure, aiming to explore its complex genetic correlation patterns with different complex traits and diseases available on BIGA. We first queried systolic blood pressure[1] from the IEU OpenGWAS (trait id: ieu-b-38, https://gwas.mrcieu.ac.uk/datasets/ieu-b-38/). BIGA performed harmonization for this queried dataset and then used the harmonized data to do LDSC massive analysis with curated datasets of European ancestry. The curated datasets included 15,428 phenotypes in total: 4,347 UK Biobank (UKB) phenotypes provided by the Neale Lab (http://www.nealelab.is/uk-biobank), 303 clinical outcomes from FinnGen, 3,905 brain imaging phenotypes from the Oxford BIG40, 3,012 brain and organ imaging phenotypes from the BIG-KP, 2,940 proteins from UKB-PPP, 897 phenotypes from GWAS Catalog, and 24 traits from PGC and other brain disorder datasets. At a false discovery rate 5% level ($P$ range = ($5.44\times10^{-244}$, $4.00\times10^{-2}$)), we found that systolic blood pressure was widely associated with different categories of phenotypes among the curated datasets (**Fig. S4**). Besides well-known associations with diastolic blood pressure and hypertension, we found 293 additional significant associations in Neale Lab, 64 in FinnGen, 265 in Oxford BIG40, 226 in BIG-KP, 74 in UKB-PPP, and 282 in GWAS Catalog (**Table S2**). For example, systolic blood pressure had significant genetic correlations with heart disease-related traits (Ischaemic heart disease, Illnesses of mother: Heart disease, Illnesses of father: Heart disease) in Neale Lab (correlation = 0.31, 0.38, and 0.29, $P = 1.05\times10^{-28}$, $2.98\times10^{-24}$, $1.36\times10^{-19}$ respectively), cardiovascular diseases in FinnGen (correlation = 0.53, $P$ = $4.41\times10^{-136}$), and coronary atherosclerosis in GWAS Catalog (correlation = 0.33, $P$= $2.21\times10^{-29}$). As expected, among all imaging traits, the strongest associations were observed on the heart imaging traits (49 were significant traits, $P < 2.30\times10^{-3}$). These traits included regional/global myocardial wall thickness, left ventricular myocardial mass, ascending/descending aorta distensibility, regional/global radial strain, and left/right ventricular cardiac output. Second, we found 437 significantly associated proteins in UKB-PPP, with the strongest genetic correlations being observed on MMP7 (correlation = 0.25, $P = 1.73\times10^{-04}$), PTPRB (correlation = 0.23, $P = 1.58\times10^{-07}$), and SIGLEC7 (correlation = 0.21, $P = 4.95\times10^{-04}$). We also performed genetic correlation analysis with diastolic blood pressure[1] (trait id: ieu-b-39, https://gwas.mrcieu.ac.uk/datasets/ieu-b-39/) and found similar association patterns to systolic blood pressure (**Fig. S5** and **Table S3**). We applied the SumHer method to repeat these analyses (**Fig. S6** and **Tables S4-S5**) and observed that the results from LDSC and SumHer were generally consistent (**Fig. S7**, Pearson's correlation = 0.9273).

To further investigate the connection between the two blood pressure measures and cardiovascular diseases, we conducted a local genetic correlation analysis with coronary artery disease[2] (CAD) through BIGA pairwise analysis using LAVA. LAVA identified 86 local genetic regions where CAD showed genetic correlations with systolic blood pressure, with 83 exhibiting positive genetic correlations and 3 showing negative correlations. Similarly, diastolic blood pressure was associated with CAD in 77 regions, 70 of which were positive (**Tables S6-S7**). We further explored the cross-population genetic correlation of blood pressure between European and East Asian ancestries. By querying East Asian diastolic and systolic blood pressure traits[3] from the GWAS Catalog (study ID: GCST90278657), we conducted Popcorn analysis using BIGA. The cross-trait genetic correlation estimates were 0.8345 for systolic and 0.7617 for diastolic blood pressure measures, neither of which were significantly different from 1 (*P* = 0.2538 and 0.1726, respectively, **Table S8**).

### Supplementary Figures

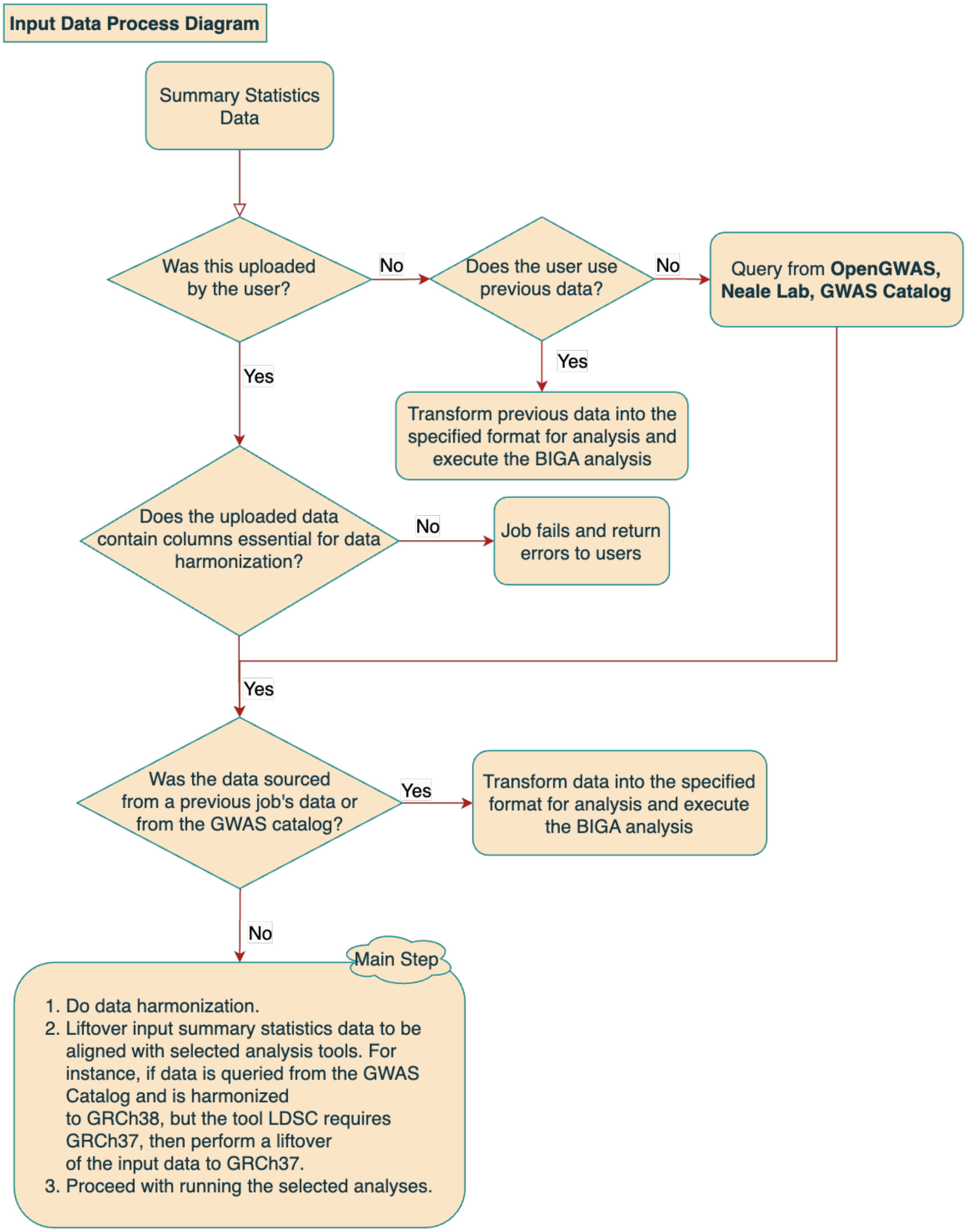


**Figure S1. Input data processing diagram.** This figure illustrates how BIGA processes different types of input data.

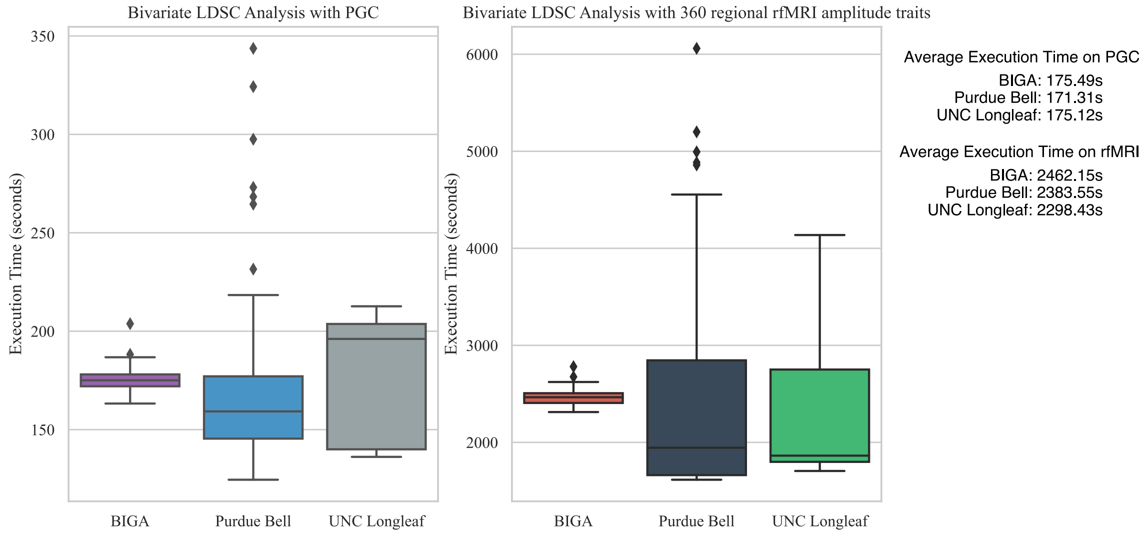


**Figure S2. Execution time on different computing platforms.** BIGA's computing server shows higher stability in terms of execution time than Purdue Bell and UNC Longleaf computing clusters.

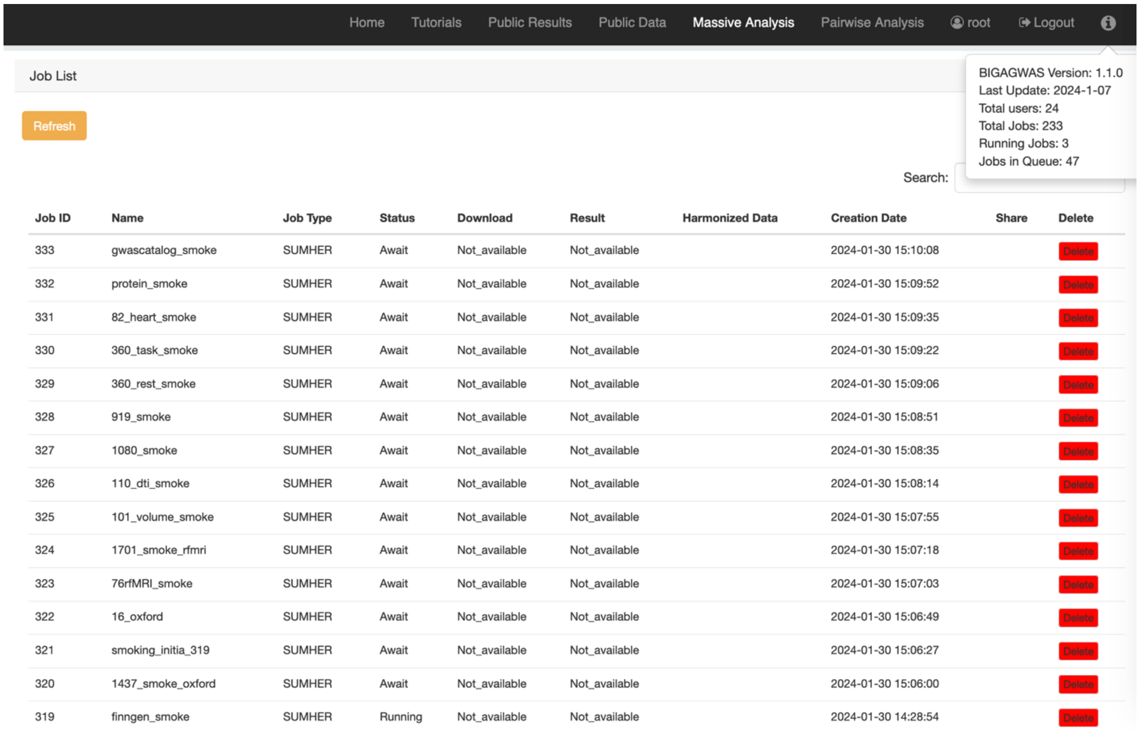

Home Tutorials Public Results Public Data Massive Analysis Pairwise Analysis root Logout

Job List

Refresh

BIGAGWAS Version: 1.1.0
Last Update: 2024-1-07
Total users: 24
Total Jobs: 233
Running Jobs: 3
Jobs in Queue: 47

Search:

| Job ID | Name | Job Type | Status | Download | Result | Harmonized Data | Creation Date | Share | Delete |
|---|---|---|---|---|---|---|---|---|---|
| 333 | gwascatalog_smoke | SUMHER | Await | Not_available | Not_available | | 2024-01-30 15:10:08 | | Delete |
| 332 | protein_smoke | SUMHER | Await | Not_available | Not_available | | 2024-01-30 15:09:52 | | Delete |
| 331 | 82_heart_smoke | SUMHER | Await | Not_available | Not_available | | 2024-01-30 15:09:35 | | Delete |
| 330 | 360_task_smoke | SUMHER | Await | Not_available | Not_available | | 2024-01-30 15:09:22 | | Delete |
| 329 | 360_rest_smoke | SUMHER | Await | Not_available | Not_available | | 2024-01-30 15:09:06 | | Delete |
| 328 | 919_smoke | SUMHER | Await | Not_available | Not_available | | 2024-01-30 15:08:51 | | Delete |
| 327 | 1080_smoke | SUMHER | Await | Not_available | Not_available | | 2024-01-30 15:08:35 | | Delete |
| 326 | 110_dti_smoke | SUMHER | Await | Not_available | Not_available | | 2024-01-30 15:08:14 | | Delete |
| 325 | 101_volume_smoke | SUMHER | Await | Not_available | Not_available | | 2024-01-30 15:07:55 | | Delete |
| 324 | 1701_smoke_rfmri | SUMHER | Await | Not_available | Not_available | | 2024-01-30 15:07:18 | | Delete |
| 323 | 76rfMRI_smoke | SUMHER | Await | Not_available | Not_available | | 2024-01-30 15:07:03 | | Delete |
| 322 | 16_oxford | SUMHER | Await | Not_available | Not_available | | 2024-01-30 15:06:49 | | Delete |
| 321 | smoking_initia_319 | SUMHER | Await | Not_available | Not_available | | 2024-01-30 15:06:27 | | Delete |
| 320 | 1437_smoke_oxford | SUMHER | Await | Not_available | Not_available | | 2024-01-30 15:06:00 | | Delete |
| 319 | finngen_smoke | SUMHER | Running | Not_available | Not_available | | 2024-01-30 14:28:54 | | Delete |

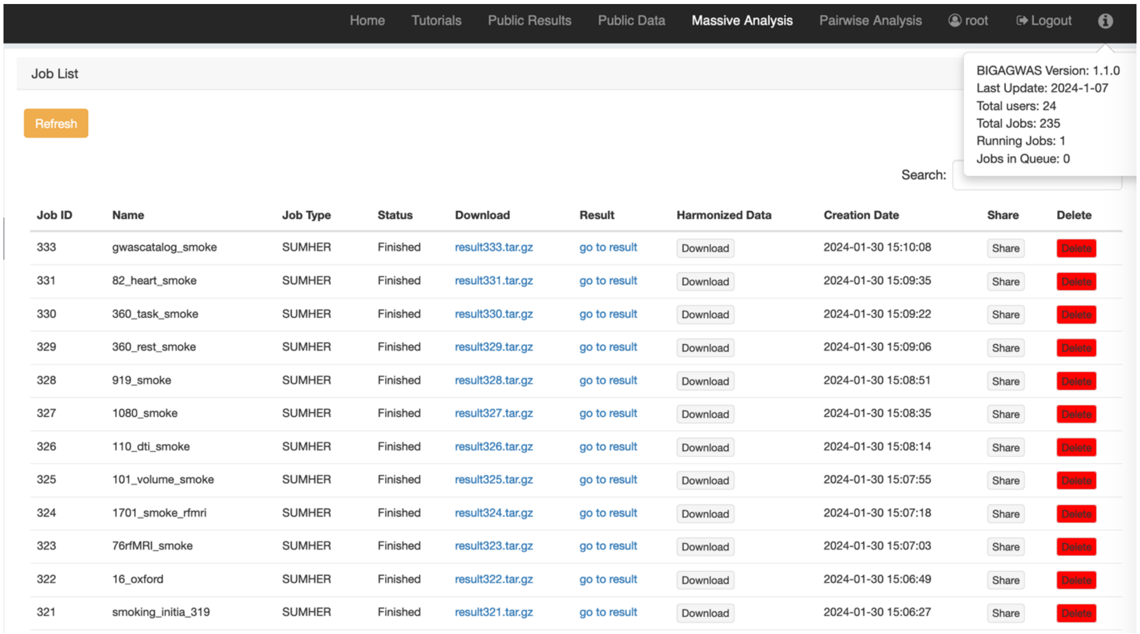

Home Tutorials Public Results Public Data Massive Analysis Pairwise Analysis root Logout

Job List

Refresh

BIGAGWAS Version: 1.1.0
Last Update: 2024-1-07
Total users: 24
Total Jobs: 235
Running Jobs: 1
Jobs in Queue: 0

Search:

| Job ID | Name | Job Type | Status | Download | Result | Harmonized Data | Creation Date | Share | Delete |
|---|---|---|---|---|---|---|---|---|---|
| 333 | gwascatalog_smoke | SUMHER | Finished | result333.tar.gz | go to result | Download | 2024-01-30 15:10:08 | Share | Delete |
| 331 | 82_heart_smoke | SUMHER | Finished | result331.tar.gz | go to result | Download | 2024-01-30 15:09:35 | Share | Delete |
| 330 | 360_task_smoke | SUMHER | Finished | result330.tar.gz | go to result | Download | 2024-01-30 15:09:22 | Share | Delete |
| 329 | 360_rest_smoke | SUMHER | Finished | result329.tar.gz | go to result | Download | 2024-01-30 15:09:06 | Share | Delete |
| 328 | 919_smoke | SUMHER | Finished | result328.tar.gz | go to result | Download | 2024-01-30 15:08:51 | Share | Delete |
| 327 | 1080_smoke | SUMHER | Finished | result327.tar.gz | go to result | Download | 2024-01-30 15:08:35 | Share | Delete |
| 326 | 110_dti_smoke | SUMHER | Finished | result326.tar.gz | go to result | Download | 2024-01-30 15:08:14 | Share | Delete |
| 325 | 101_volume_smoke | SUMHER | Finished | result325.tar.gz | go to result | Download | 2024-01-30 15:07:55 | Share | Delete |
| 324 | 1701_smoke_rfmri | SUMHER | Finished | result324.tar.gz | go to result | Download | 2024-01-30 15:07:18 | Share | Delete |
| 323 | 76rfMRI_smoke | SUMHER | Finished | result323.tar.gz | go to result | Download | 2024-01-30 15:07:03 | Share | Delete |
| 322 | 16_oxford | SUMHER | Finished | result322.tar.gz | go to result | Download | 2024-01-30 15:06:49 | Share | Delete |
| 321 | smoking_initia_319 | SUMHER | Finished | result321.tar.gz | go to result | Download | 2024-01-30 15:06:27 | Share | Delete |

**Figure S3. Submission, querying, and completion of 50 jobs.**

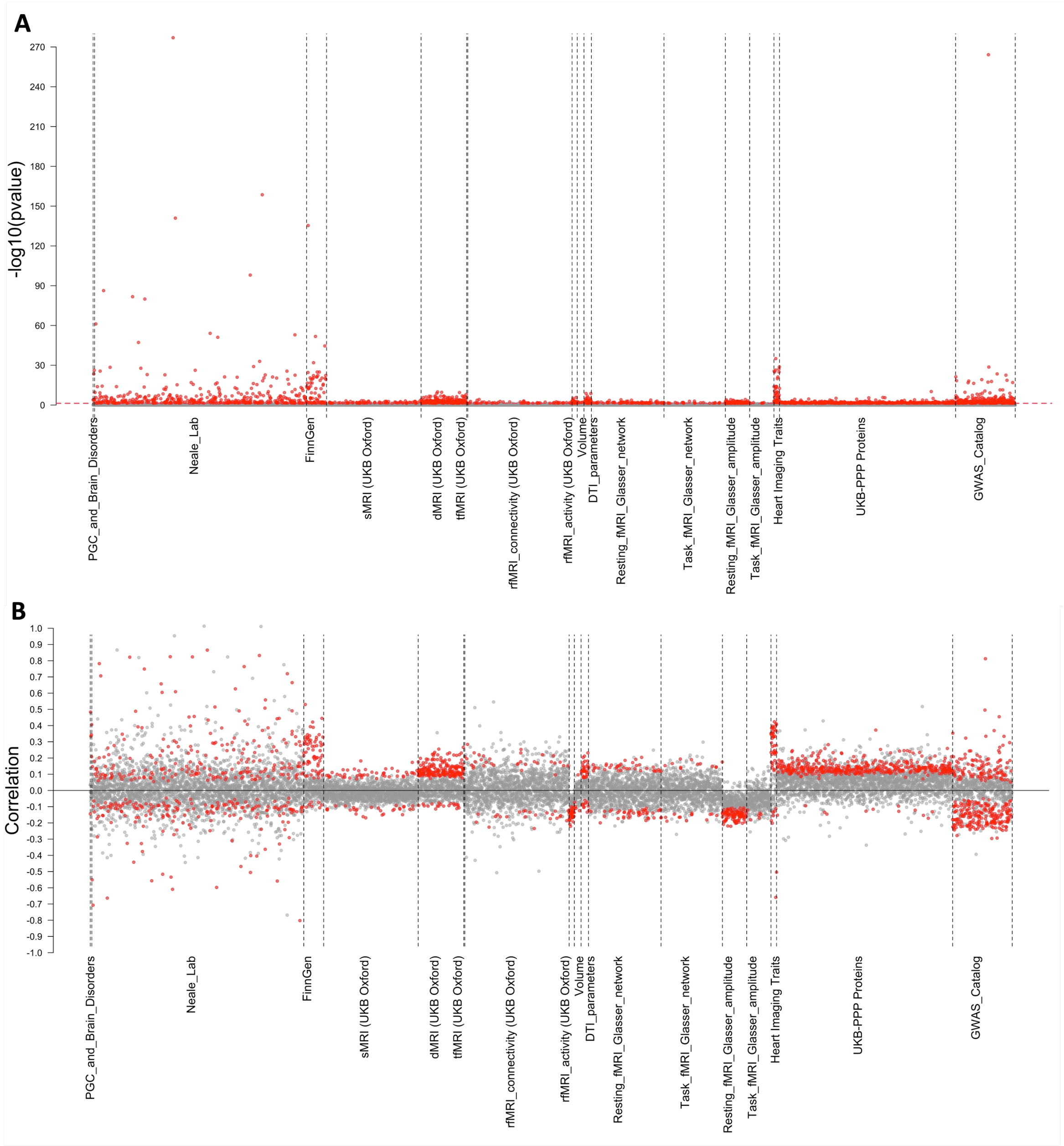


**Figure S4. Genetic correlations between systolic blood pressure and all European ancestry curated datasets in LDSC analysis. (A)** and **(B)** show the *P* values and genetic correlation estimates, respectively. Significant correlations at a false discovery rate 5% level are highlighted in red color.

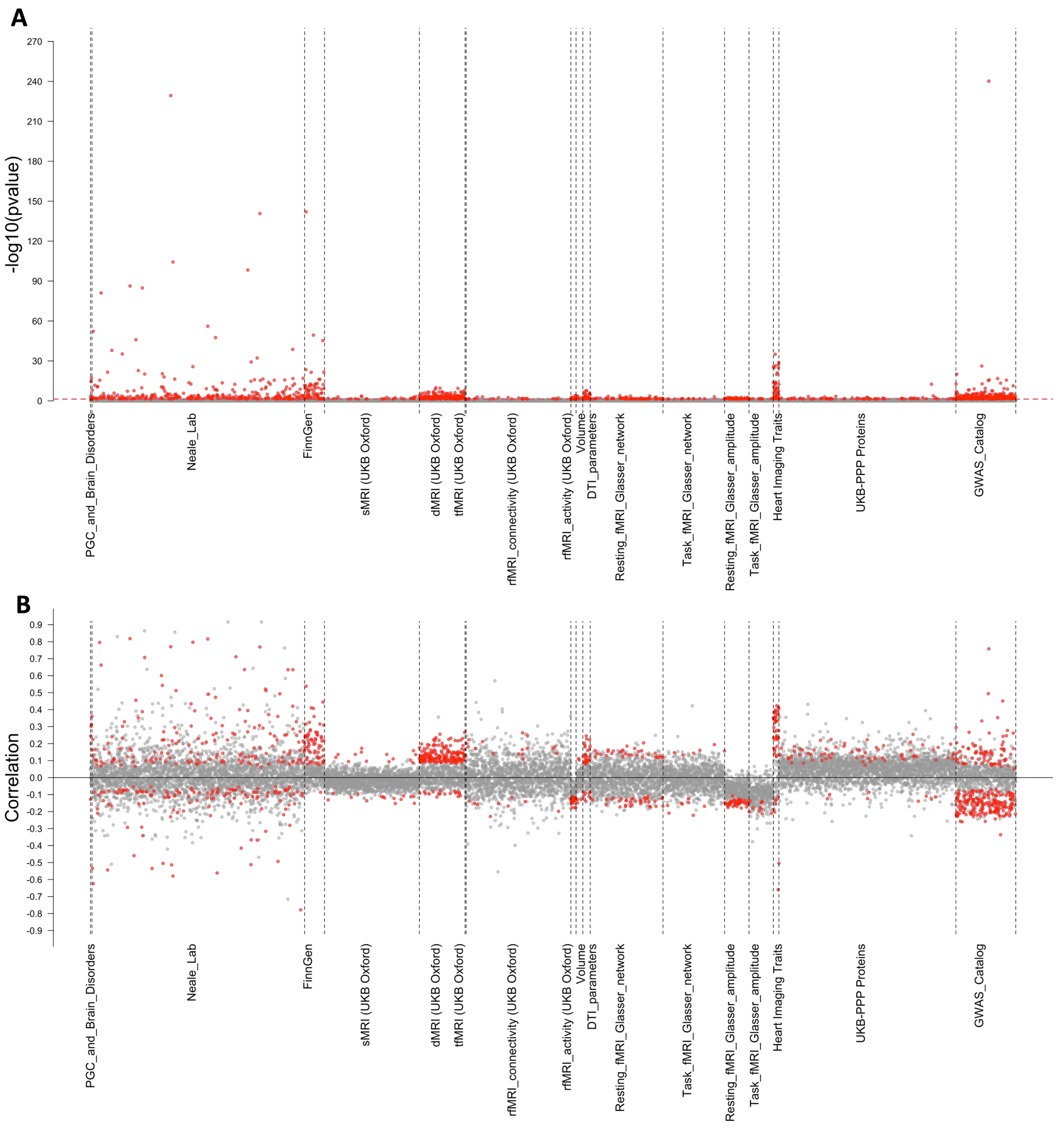


**Figure S5. Genetic correlations between diastolic blood pressure and all European ancestry curated datasets in LDSC analysis. (A)** and **(B)** show the *P* values and genetic correlation estimates, respectively. Significant correlations at a false discovery rate 5% level are highlighted in red color.

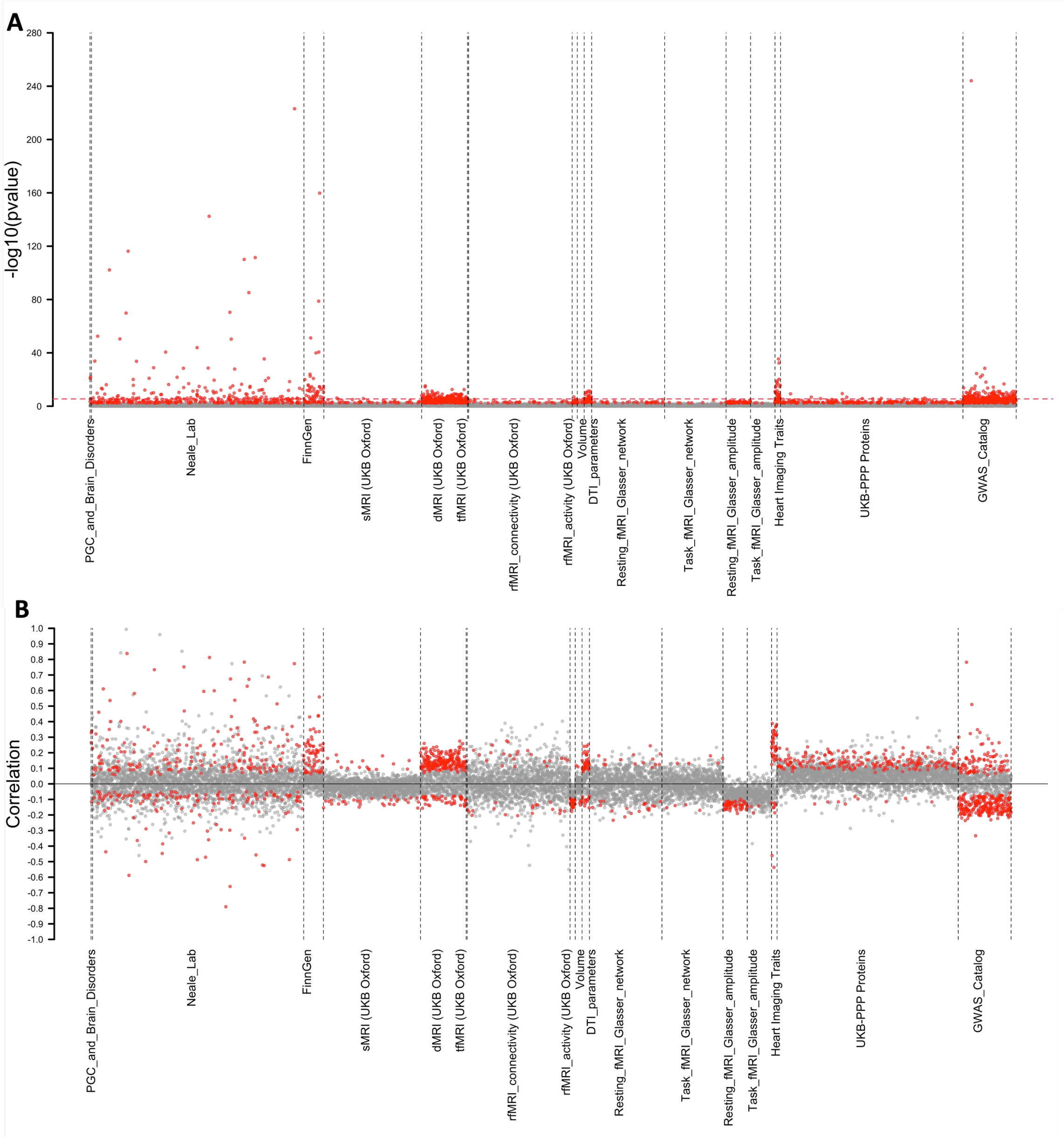


**Figure S6. Genetic correlations between diastolic blood pressure and all European ancestry curated datasets in SumHer analysis. (A)** and **(B)** show the *P* values and genetic correlation estimates, respectively. Significant correlations at a false discovery rate 5% level are highlighted in red color.

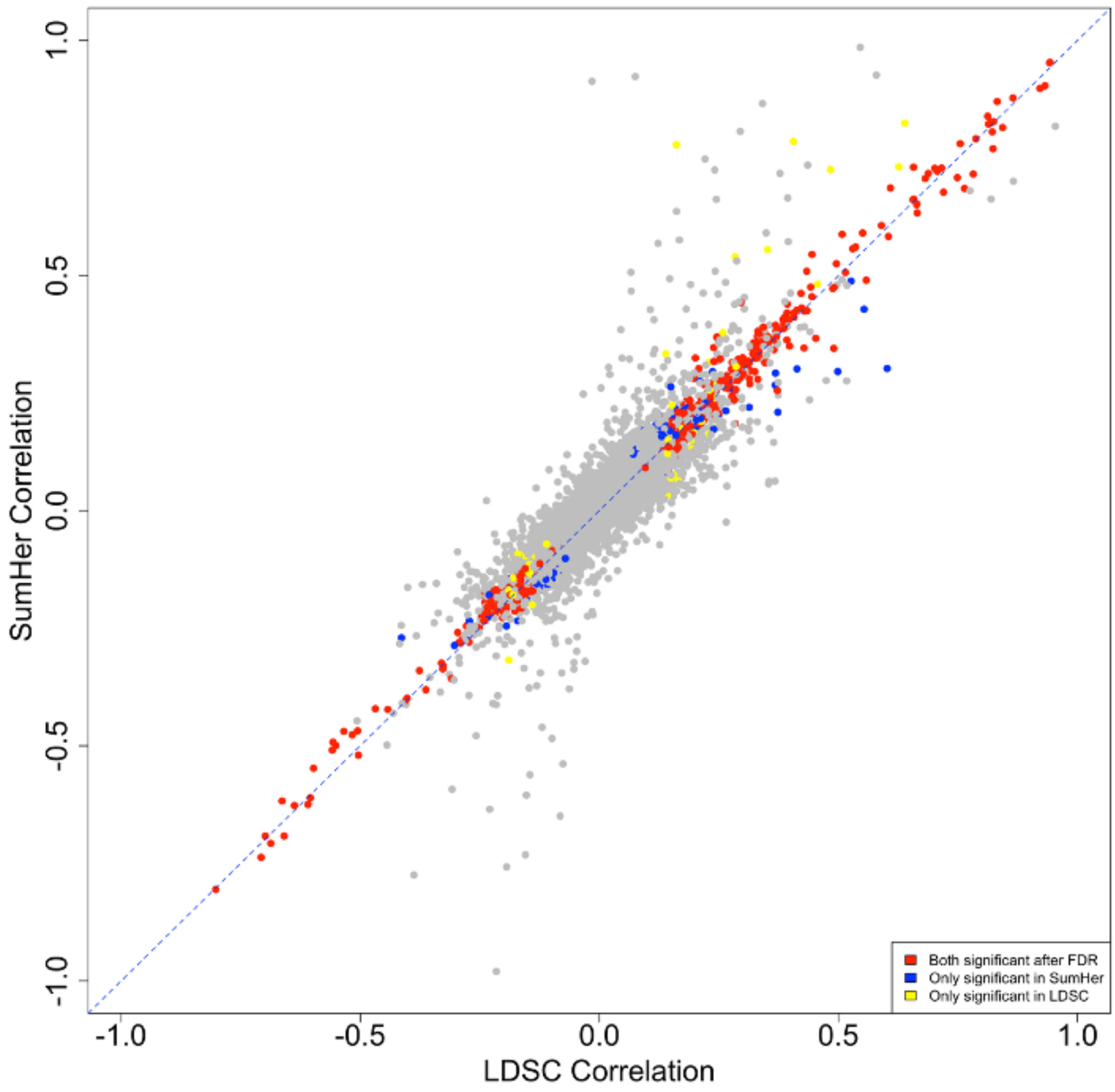


**Figure S7. Comparison of LDSC and SumHer genetic correlation estimates for systolic blood pressure.** We highlight traits significant in both methods as well as those significant in only one of the two methods using distinct colors.

### **Legends for Tables S1 to S8** (All tables can be found in a zip file).

**Table S1. Column requirements for BIGA data harmonization and analysis.**

**Table S2. LDSC genetic correlations between systolic blood pressure and traits in curated datasets on BIGA.** We remove invalid genetic correlation estimates and *P* values.

**Table S3. LDSC genetic correlations between diastolic blood pressure and traits in curated datasets on BIGA.** We remove invalid genetic correlation estimates and *P* values.

**Table S4. SumHer genetic correlations between systolic blood pressure and traits in curated datasets on BIGA.** We remove invalid genetic correlation estimates and *P* values.

**Table S5. SumHer genetic correlations between diastolic blood pressure and traits in curated datasets on BIGA.** We remove invalid genetic correlation estimates and *P* values.

**Table S6. Local genetic correlations between systolic blood pressure and coronary artery disease.**

**Table S7. Local genetic correlations between diastolic blood pressure and coronary artery disease.**

**Table S8. Popcorn cross-ancestry genetic correlations between East Asian and European blood pressure measures.** We test for the hypothesis that the genetic correlation is 1.
